# Activation Mechanism of Strigolactone Receptors And Its Impact On Ligand Selectivity Between Host And Parasitic Plants

**DOI:** 10.1101/2021.06.02.446815

**Authors:** Jiming Chen, David C. Nelson, Diwakar Shukla

## Abstract

Parastic weeds such as *Striga* have led to significant losses in agricultural productivity worldwide. These weeds use the plant hormone strigolactone as a germination stimulant. Strigolactone signaling involves substrate binding and hydrolysis followed by a large conformational change of the receptor to a “closed” or “active” state that is able to associate with a downstream signaling partner MAX2/D3. The crystal structure of the active and inactive *At*D14 receptor have helped in elucidating the structural changes involved in activation. However, the mechanism by which the receptor activates remains unknown. The ligand dependence of *At*D14 activation has been disputed by mutagenesis studies showing that enzymatically inactive receptors are able to form a complex with MAX2 proteins. Furthermore, activation differences between strigolactone receptor in Striga, *Sh*HTL7 and textitAtD14 could contribute to the high sensitivity to strigolactones exhibited by parasitic plants. Using molecular dynamics simulations, we demonstrate that both *At*D14 and *Sh*HTL7 could adopt an active conformation in absence of ligand. However, the *Sh*HTL7 receptor exhibits higher population in the inactive *apo* state as compared to the *At*D14 receptor. We demonstrate that this difference in inactive state population is caused by sequence differences between their D-loops and its interactions with the catalytic histidine that prevents full binding pocket closure in *Sh*HTL7. These results indicate that hydrolysis of a strigolactone ligand would enhance the active state population by destabilizing the inactive state in *Sh*HTL7 as compared to *At*D14. We also show that the mechanism of activation is more concerted in *At*D14 than in *Sh*HTL7 and that the main barrier to activation in *Sh*HTL7 is closing of the binding pocket.

## 1 Introduction

Strigolactones are a class of endogenous plant hormones responsible for regulating shoot branching and root architecture in plants^1–4^. They also function as a germination stimulant for members of the *Striga* genus of parasitic weeds^5^. The strigolactone signaling process is known to involve enzymatic hydrolysis of a strigolactone molecule and a large conformational change of the receptor to an “active” state that allows it to associate with its signalling partner, MAX2^6, 7^. The strigolactone receptor also associates with SMXL proteins which are then ubiquitinated and degraded^6,8,9^. The active state of the *At*D14 receptor was crystallized in complex with *Os*D3, the rice ortholog of *At*MAX2^6^. While crystal structures have played a critical role in elucidating the active structure of D14 which associates with its signaling partner, a full mechanistic picture of the process by which D14/HTL proteins undergo the transition between their inactive and active states remains unknown.

The “active” state crystal structure showed the *At*D14 receptor in complex with a covalently linked intermediate (CLIM), which is thought to be an intermediate species of strigolactone hydrolysis. A covalently bound hydrolysis intermediate has also been reported via mass spectrometry as well^10^. A reanalysis of this crystal structure by Carlsson *et al*. attributed the electron density of the CLIM molecule to an iodide ion^11^. However, another reanalysis of this structure suggested that the present species is a covalently bound D-ring as suggested by de Saint Germain *et al*.^10, 12^. Additionally, Seto *et al*. found that while mutation of any of the catalytic triad residues (S97, D218, H247) abolishes their enzymatic activity, the D218A mutant alone is still able to complement the *atd14* mutant phenotype. This suggests that the catalytically inactive D218A mutant of *At*D14 is able to transduce a SL signal, which implies that SL hydrolysis is not strictly necessary for receptor activation^7^. It is unclear how the D218A catalytic triad mutation leads to constitutive receptor activity in *At*D14.

Despite high sequence, structure, and binding site conservation, one receptor in *Striga, Sh*HTL7, uniquely confers germination responses to picomolar GR24, a synthetic strigolactone analog, when expressed in *Arabidopsis thaliana*.^13^. Since this sensitivity to the substrate is measured in terms of downstream signaling, it is possible that receptor activation plays a role in modulating this difference in sensitivity to GR24. However, the mechanism by which the activation process modulates ligand sensitivity is unknown. Recently, Wang *et al*. determined that strong association with the MAX2 signaling partner distinguishes *Sh*HTL7 from other *Sh*HTL proteins^14^. One possible explanation for this enhanced association is higher stability of the active, i.e. association-capable conformation of the protein in *Sh*HTL7.

A powerful method of elucidating mechanistic details of this activation process is long timescale molecular dynamics (MD) simulations. MD simulations have been widely employed to resolve atomic level details about the activation process of signaling proteins relevant to human health^16–20^. Recently, MD simulations have been used to study conformational changes associated with substrate binding in plant hormone receptors and have provided insights into the molecular determinants of hormone affinity differences between receptors and regulation of the receptor activation process.^21–29^. Despite the importance of strigolactone perception in parasitic plants and the availability of structural informaiton, MD simulations have not been used to characterize the full activation process of strigolactone receptors in plants.

Recently, Lee *et al*. used MD simulations combined with mutagenesis experiments to determine that mutants that increase the flexibility of DAD2, the petunia ortholog of *At*D14, are able to enhance its interaction with its MAX2 signaling partner^30^. While this provides an important insight into regions of the protein that could mediate its ability to activate, the information provided by the MD simulations is somewhat limited since the transition from the inactive to active states of the receptor was not captured in full. In our prior work^31^, we have used long timescale MD simulations to investigate the ligand binding mechanism for strigolactone receptors and its potential impact on the ligand selectivity exhibited by receptors in parasitic plants. This work uncovered several factors that could enhance SL binding and catalytic activity in *Sh*HTL7 in relation to *At*D14. Briefly, *Sh*HTL7 may be more efficient at binding SL in a catalytically productive conformation, while additional flexibility of the *At*D14 binding pocket enables the ligand to bind in unproductive conformations. However, further work is needed to elucidate the mechanism of subsequent steps of strigolactone signaling and differences between species. The downstream strigolactone signaling is regulated by ligand binding, hydrolysis, receptor activation, association with MAX2 proteins and SMXL proteins and their degradation. Therefore, a comprehensive investigation of these processes is needed to establish their relative importance in regulating strigolactone signaling in both host and parasitic plants.

Here, we utilize long-timescale (∼600 *µ*s aggregate) unbiased MD simulations which provide a high-resolution dynamical view of *At*D14 and *Sh*HTL7 activation in their *apo* (ligand-free)forms. Markov State Models (MSMs) built from the simulation data were used to estimate the thermodynamics and kinetic properties associated with the activation process. We demonstrate that the activation process for strigolactone receptors is controlled by a complex network of molecular or conformational switches. The main barriers to receptor activation are detachment of the D-loop and closure of the binding pocket via the T1 and T3 helices. We also identify the intermediate states that connect the inactive and active states of strigolactone receptors and could be targeted for inhibitor design. Additionally, we identify the several key residue interactions with H247 that are responsible for differences in the activation mechanisms of *At*D14 and *Sh*HTL7.

## 2 Results

### Transition pathways from inactive to active states

Strigolactone receptor activation involves four molecular or conformational switches^6, 32, 33^: (1) closure of the binding pocket via merging of the T1 and T3 helices, (2) partial unfolding of the T2 helix, (3) extension of the T1 helix by the folded portion of the T2 helix, and (4) detachment of the D-loop. To gain quantitative insight into the activation process, we first computed free energy landscapes of the activation processes of both *At*D14 and *Sh*HTL7 projected onto four metrics associated with each of the four molecular switches (Fig. 2). Briefly, these landscapes were obtained by calculating a probability density along each pair of metrics and using the relation *F* = -*RT* ln(*P*) to convert the probability densities to free energies. From projection of our data onto each of these molecular switches (Fig. 2), we can see that both *At*D14 and *Sh*HTL7 are able to undergo activation as *apo* proteins. However, the dominant pathways that connect the active and inactive states in the two proteins are different. Notably, a set of intermediate states observed in *At*D14 in which the T1-T3 distance and T2 helical content are low. However, these intermediates are less present in *Sh*HTL7. This is an indication that there is less coupling between these two molecular switches in *Sh*HTL7 than in *At*D14.

**Fig. 1.**
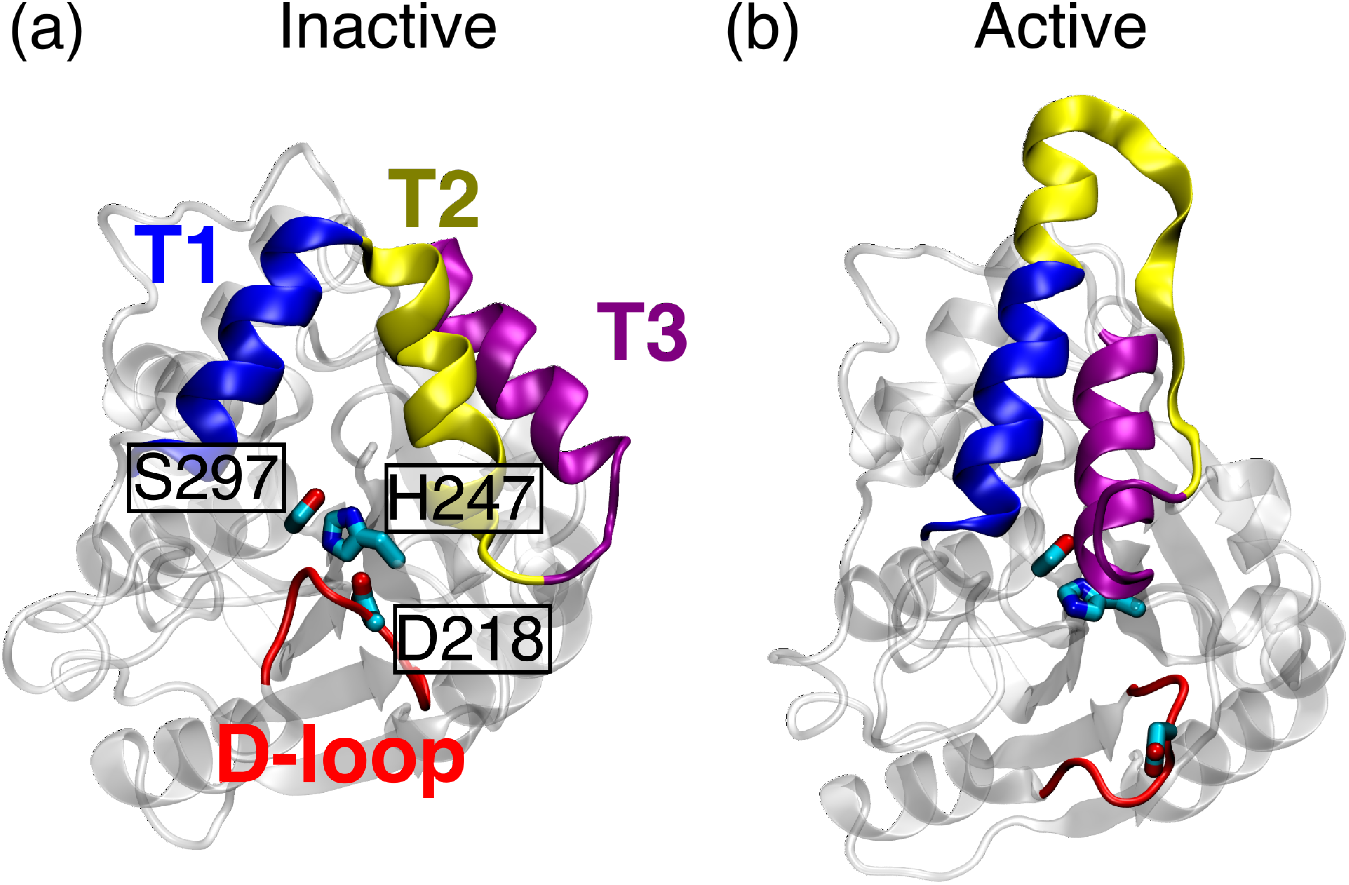
Inactive (a) and active (b) states of *At*D14. The T1, T2, and T3, helices are shown in blue, yellow, and purple, respectively, and the aspartate loop (D-loop) is shown in red. The inactive state was obtained from PDB ID 4IH4^15^, and the active state was obtained from PDB ID 5HZG^6^. Missing D-loop residues in the active state were added using Modeller.

**Fig. 2.**
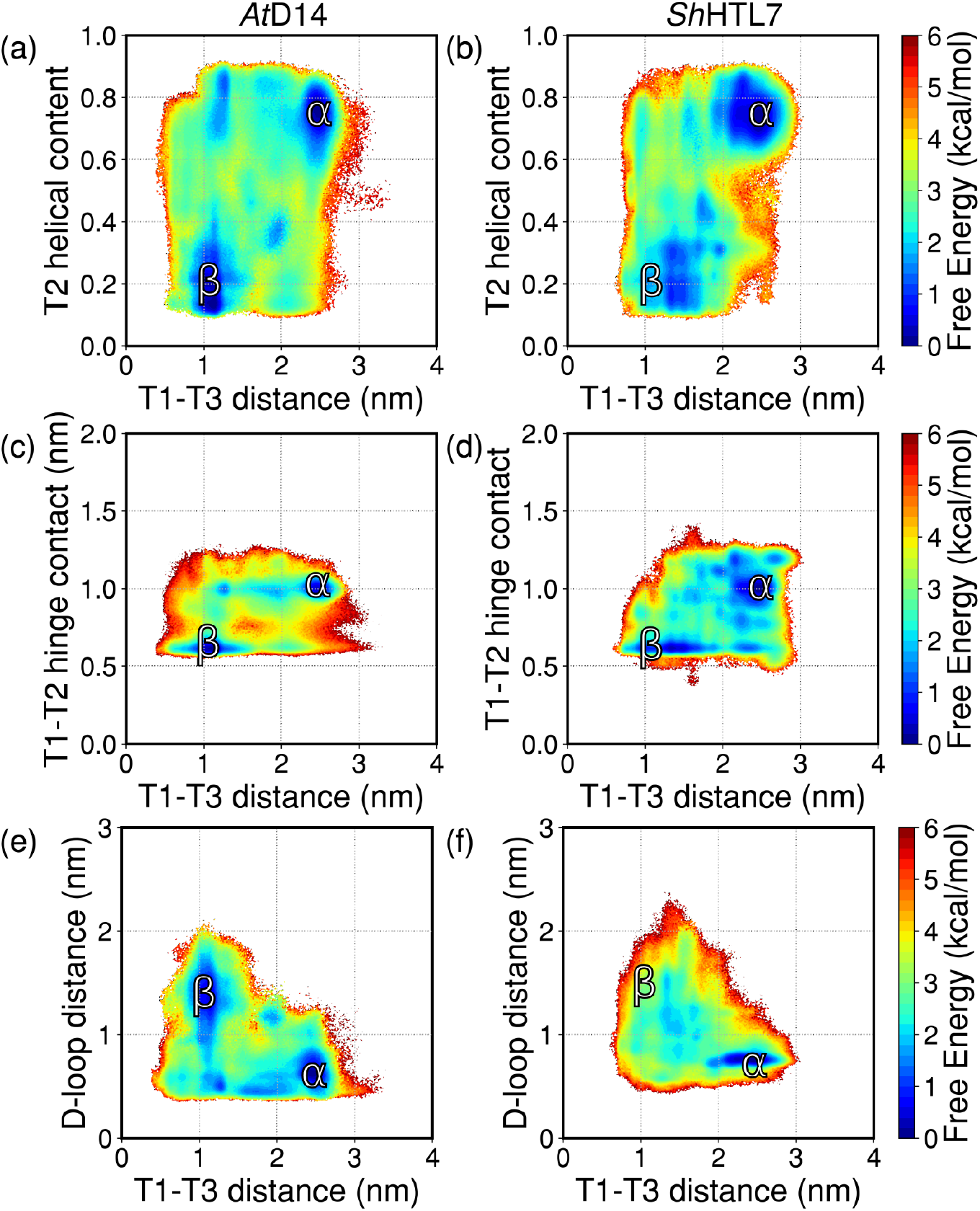
Free energy landscapes of the activation process in *At*D14 (a,c,e) and *Sh*HTL7 (b,d,f). These landscapes show the activation process in terms of pocket closing by merging of the T1 and T3 helices and unfolding of the T2 helix (a,b), extension of the T1 helix by the folded portion of the T2 helix (c,d), and D-loop detachment (e,f). The active state in each landscape is labeled as *α*, and the inactive state is labeled as *β*.

### Active state is more stable in *At*D14 than in *Sh*HTL7

Our free energy landscapes indicate that both *At*D14 and *Sh*HTL7 are able to undergo activation in their *apo* states. However, the is higher in *At*D14 than in *Sh*HTL7. While the active and inactive states have similar populations as indicated by their free energies in *At*D14, the free energy of the active state in *Sh*HTL7 is ∼2-3 kcal/mol higher than that of the inactive state. This indicates that *Sh*HTL7 is roughly 3-5 times more likely to occupy its inactive state than its active state. For both *At*D14 and *Sh*HTL7, the free energy barrier for transitioning between the inactive to active state is ∼3 kcal/mol. To evaluate this barrier, we performed time-lagged independent component analysis (TICA)^34^. TICA coordinates are linear combinations of structural metrics that correspond to the slowest processes captured in simulations. The slowest processes are the processes associated with the highest free energy barriers, so computing free energy landscapes on TICA coordinates gives a more accurate estimate of free energies associated with the activation process. Free energy landscapes of the activation processes projected onto these slow coordinates are shown in Fig. S1. To compare the slow processes in *At*D14 and *Sh*HTL7, we projected the simulation data of each protein onto the TICA coordinates of the other protein. Cross-projection of each dataset onto each set of TICA coordinates additionally indicates that although both *At*D14 and *Sh*HTL7 are able to access both the inactive and active states, the transition pathways between the inactive and active states are different in the two proteins. This indicates that the main barriers to transitioning from the inactive to active states are different.

### Coupled activation of molecular switches regulates the conformational dynamics of strigolactone receptors

To investigate the differences in the activation mechanism between *At*D14 and *Sh*HTL7, we computed conditional probabilities of each of the molecular switches to turn “on” or become “active” given that other three molecular switches are in the “on” state as well. Table 1 shows the conditional and the equilibrium probabilities for each individual molecular switch to be in the “on” state. The equilibrium probabilities of each of the individual molecular switches are close to ∼50% in *At*D14 and between ∼10-30% in *Sh*HTL7, which is consistent with the high active state population observed for *Sh*HTL7 in the free energy landscapes. The table also indicates that the dynamics of the molecular switches are coupled - the activation of any one molecular switch increases the probabilities of all of the others switches to activate. In particular, the probability of the T2 helix unfolding increases to *>*80% from ∼54% and ∼32% in both *At*D14 and *Sh*HTL7, respectively. Similarly, extension of T1 and closure of the binding pocket via the T1 and T3 helices are also highly coupled molecular switches. In *At*D14, the probabilities of both of these metrics rises to *>*87% with other switches in “on” state. In *Sh*HTL7, the probability of T1 extension rises to ∼89% when the T1 and T3 helices are close together, however, the probability of the T1 and T3 helices being close together only rises to ∼31% when the T1 helix is extended. This low probability indicates that a higher barrier to the merging of the T1 and T3 helices is likely a basis for the stability difference in the active state population of *At*D14 and *Sh*HTL7. Finally, while the probability of D-loop detachment increases with the presence of any other active switches, the probabilities of D-loop detachment are lower in *Sh*HTL7 (∼55-63%) than in *At*D14 (∼77-84%). This also indicates that a difference in the probability of D-loop detachment is a differentiating factor between the activation processes of *At*D14 and *Sh*HTL7.

**Table 1.**
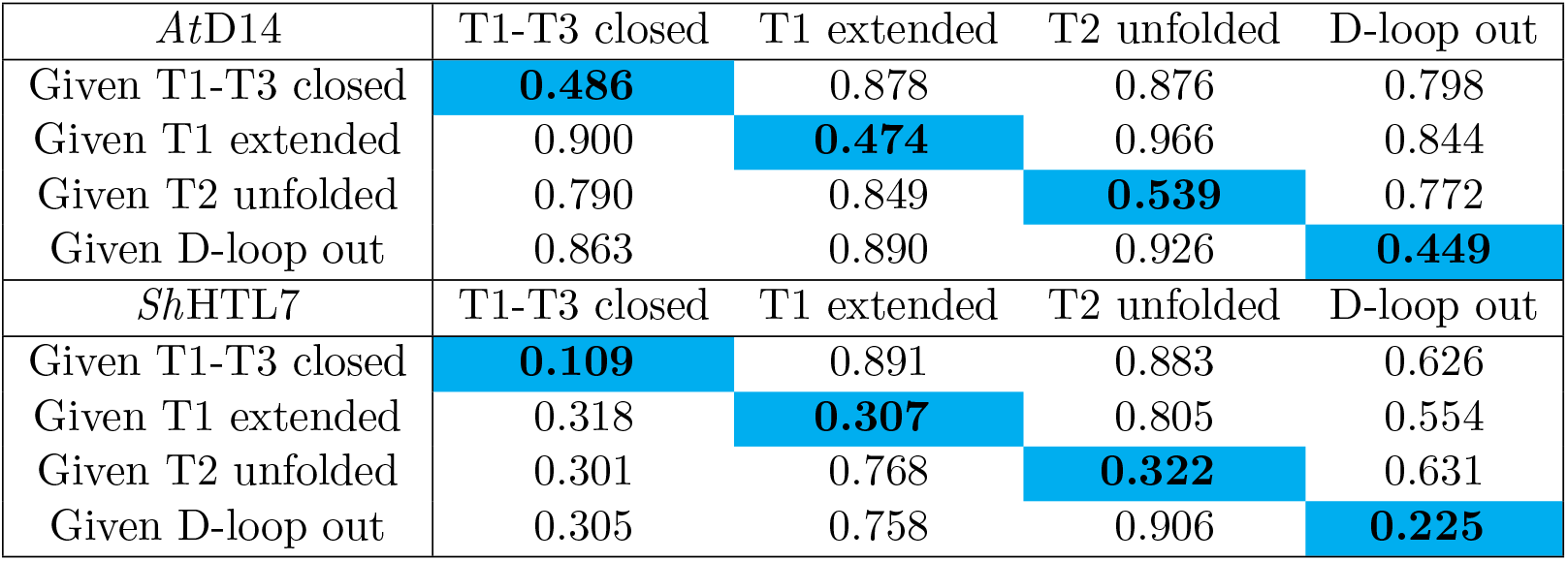
Conditional probabilities for each molecular switch given the presence of each other molecular switch in *apo At*D14 and *Sh*HTL7. Each column of the table indicates the probabilities for each molecular switch, and each row corresponds to a different condition. Highlighted values along the diagonal are equilibrium probabilities for each individual molecular switch.

In addition to computing conditional probabilities of each molecular switch given each of the other molecular switch are individually “on”, we also computed the conditional probabilities of each molecular switch given the presence of each pair of molecular switches (Table S1) and given the presence of all other molecular switches (Table 2). In *At*D14, these values indicate that the activation is a highly concerted process - when any three molecular switches are present, the remaining molecular switch has a ∼87-99% chance of being present. In *Sh*HTL7, the presence of any three molecular switches increases the the probability of the remaining molecular switch to *>*90%. However, the probabilities of T1-T3 closure and D-loop detachment remain much lower than the other markers, at ∼33% and ∼64%, respectively. This further indicates that differences in the closure of the binding pocket and D-loop detachment between *At*D14 and *Sh*HTL7 are modulators of the observed difference in their active state population.

**Table 2.** Conditional probabilities for each of the four molecular switches given that all other molecular switches are “on” in textitAtD14 and *Sh*HTL7.

|  | *AtD14* | *ShHTL7* |
| --- | --- | --- |
| T1-T3 closed | 0.934 | 0.356 |
| T1 extended | 0.992 | 0.929 |
| T2 unfolded | 0.985 | 0.934 |
| D-loop out | 0.877 | 0.648 |

### Effect of D-loop sequence differences on detaching motion

Previously, we identified three key residues (A216S, K217N, and S220M) in the sequence of the D-loop that stabilize the D-loop “outward” conformation in *At*D14 as compared to the *Sh*HTL7.^31^ These differences allow the receptor to be more enzymatically competent in *Sh*HTL7 than in *At*D14 due to contact between the aspartate in the D-loop and the catalytic histidine residue while the D-loop is in its “attached” or “inward” conformation. However, this has a opposing effect on the receptor activation in *apo* state. Since one of the molecular switch involved in receptor activation is detachment of the D-loop, stabilization of the D-loop “outward” or “detached” conformation is expected to enhance activation of the protein. To determine the effects of the three previously identified mutations on D-loop conformation during the receptor activation process, we computed free energy landscapes projected onto each of these contacts against D218/217-H247/246 distance (Fig. 3).

**Fig. 3.**
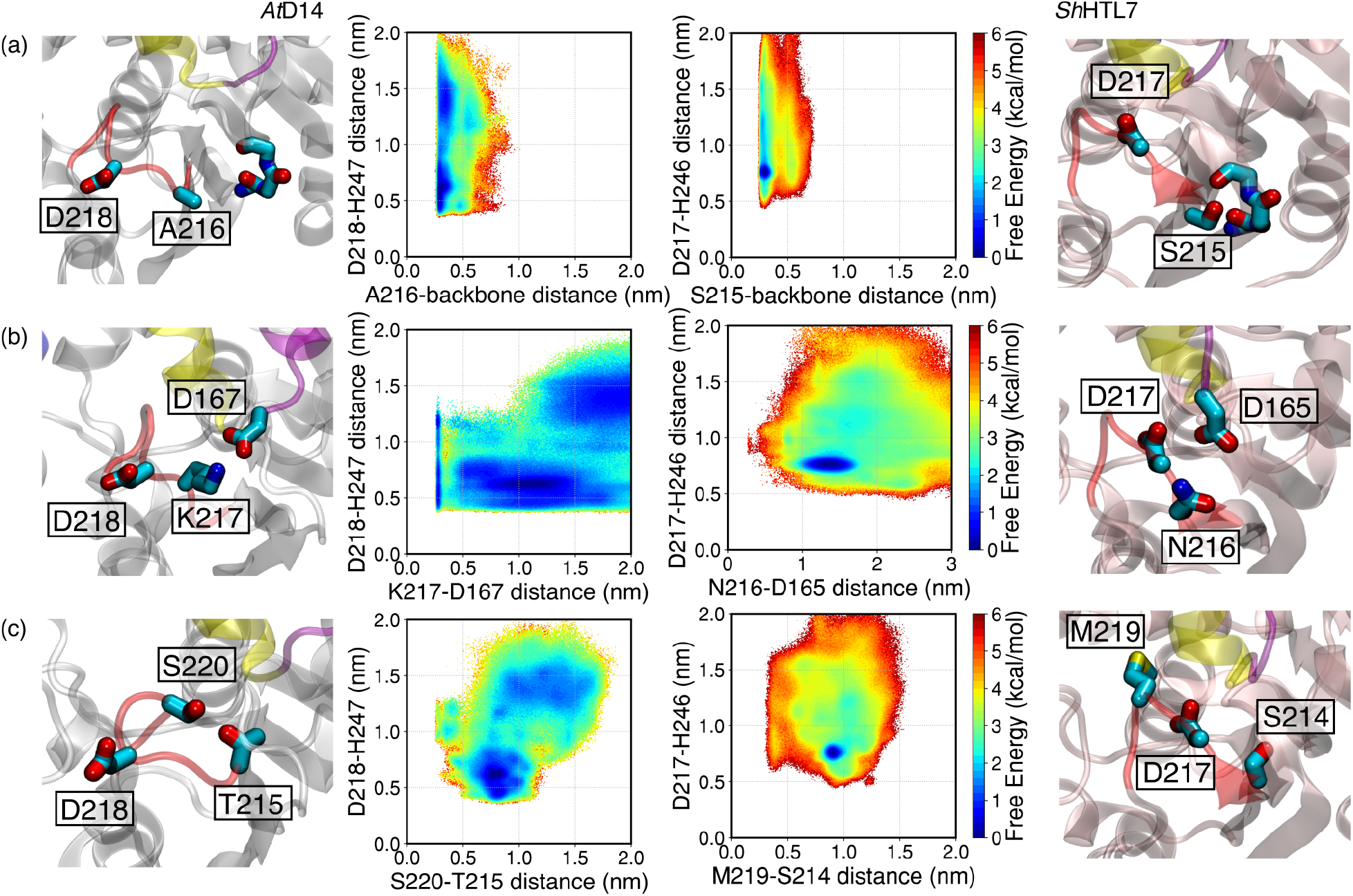
Free energy landscapes of (a) A216/S215-backbone contact distance, (b) K217/N216-D167/165 contact distance (c) S220/M219-T215/S214 contact distance projected against D218/217-H247/246 contact distance in *At*D14 and *Sh*HTL7. These interactions were previously identified to contribute to stabilization of the protein in catalytically competent, more inactive-like states in *Sh*HTL7 and catalytically incompetent, less inactive-like states in *At*D14.

The A216S substitution in *Sh*HTL7 (Fig. 3a) stabilizes the D-loop “inward” conformation, which is enzymatically competent but less active-like. In *Sh*HTL7, S215 forms a contact with neighboring backbone atoms, which reduces the range of motion of the D-loop. In *At*D14, the equivalent residue is A216, which is unable to form the same hydrogen bond with neighboring backbone atoms. As shown in the associated free energy landscapes, formation of the S215-backbone contact is highly favorable, and when this contact is formed, the D217-H246 distance is small (<8 Å), indicating that the D-loop is in its “inward” conformation. In *At*D14, however, the free energy landscape shows that the D-loop can readily adopt both “inward” and “outward” conformations. This indicates that lack of the S215-backbone hydrogen bond in *At*D14 facilitates the detachment of the D-loop. Thus, there is a lower barrier for D-loop detachment in *At*D14. An analysis of sequence conservation at this site among close homologs of *At*D14, *Sh*HTL7, and *At*KAI2 shows high variability at the site in all three groups (Fig. S5).

The K217 residue enables the formation of a salt bridge with D167 in *At*D14 that is not present in *Sh*HTL7 due to K217 N subsitution (Fig. 3b). Formation of the K217-D167 salt bridge stabilizes conformations in which D218 and H247 are far apart, thus stabilizing the D-loop “outward” conformation. In contrast, this salt bridge is absent in *Sh*HTL7 due to mutation to an uncharged residue. Thus, the interaction between N216 and D165 is weakened and unable to stabilize D-loop “outward” conformations. However, an important caveat is that D-loop “outward” conformations also have a higher relative population in *At*D14 than in *Sh*HTL7 when the K217-D167 salt bridge is not formed. This means that this salt bridge is not the strongest contributor to the more stable D-loop “outward” conformations in *At*D14. Sequences of homologs at this site show high variability among *At*D14 homologs, while the site is a highly conserved lysine among both *Sh*HTL7 and *At*KAI2 homologs (Fig. S5).

Finally, the S220 residue in *At*D14 locks the D-loop in an “outward” conformation by forming a hydrogen bond with T215 on the other end of the D-loop. The equivalent D-loop residues in *Sh*HTL7 are M219 and S214, which are unable to form a hydrogen bond, thus the relative population of conformations in which they are in contact is reduced compared to *At*D14. This allows for higher stability of D-loop “inward” conformations which are more catalytically competent but less active-like. As with K217N substitution, it is important to note that the D-loop “outward” conformations are more stable in *At*D14 than in *Sh*HTL7 even in the absence of the S220-T215 hydrogen bond. Thus, this interaction is not the strongest contributor to the difference in D-loop conformation stabilities during the activation process. Sequences of homologs at this site show high variability among *At*D14 homologs, with the majority of sequences containing a hydrophobic residue at this site. The majority of both *Sh*HTL7 and *At*KAI2 homologs contain an alanine at this site (Fig. S5).

### H247 contacts modulate differences in D-loop detachment and T1-T3 closure

To determine the sequence differences that contribute to the difference in the active state population between *Sh*HTL7 and *At*D14, we computed contact probability matrices of each pair of contacts from kinetic Monte Carlo trajectories generated from our Markov state models. Briefly, these trajectories are generated by using the MSM transition matrix to propagate the state transition of a protein from an initial state. In this case, an inactive state cluster from the MSM was used as an initial state. Full details of this procedure can be found in Methods section. Using these trajectories, we computed the equilibrium contact probabilities for each contact pair in the protein (Fig. 4). From these contact maps, a cluster of contacts around the H247 residue is identified that has different probabilities in the two receptors. Since covalent modification of H247 by a hydrolyzed ligand has been hypothesized to enhance activation in D14 proteins^6, 10^, we computed contact probabilities of each residue with H247. We then additionally computed contact probabilities of each residue with residues that have a high difference in H247 contact probability Figure S4. To identify the contacts that contribute to the differences in the pocket closing T1-T3 motion and D-loop detachment, we computed the free energy landscapes of each MD data set projected onto these contacts against T1-T3 helix distance and D-loop-H247 distance (Fig. 5, 6).

**Fig. 4.**
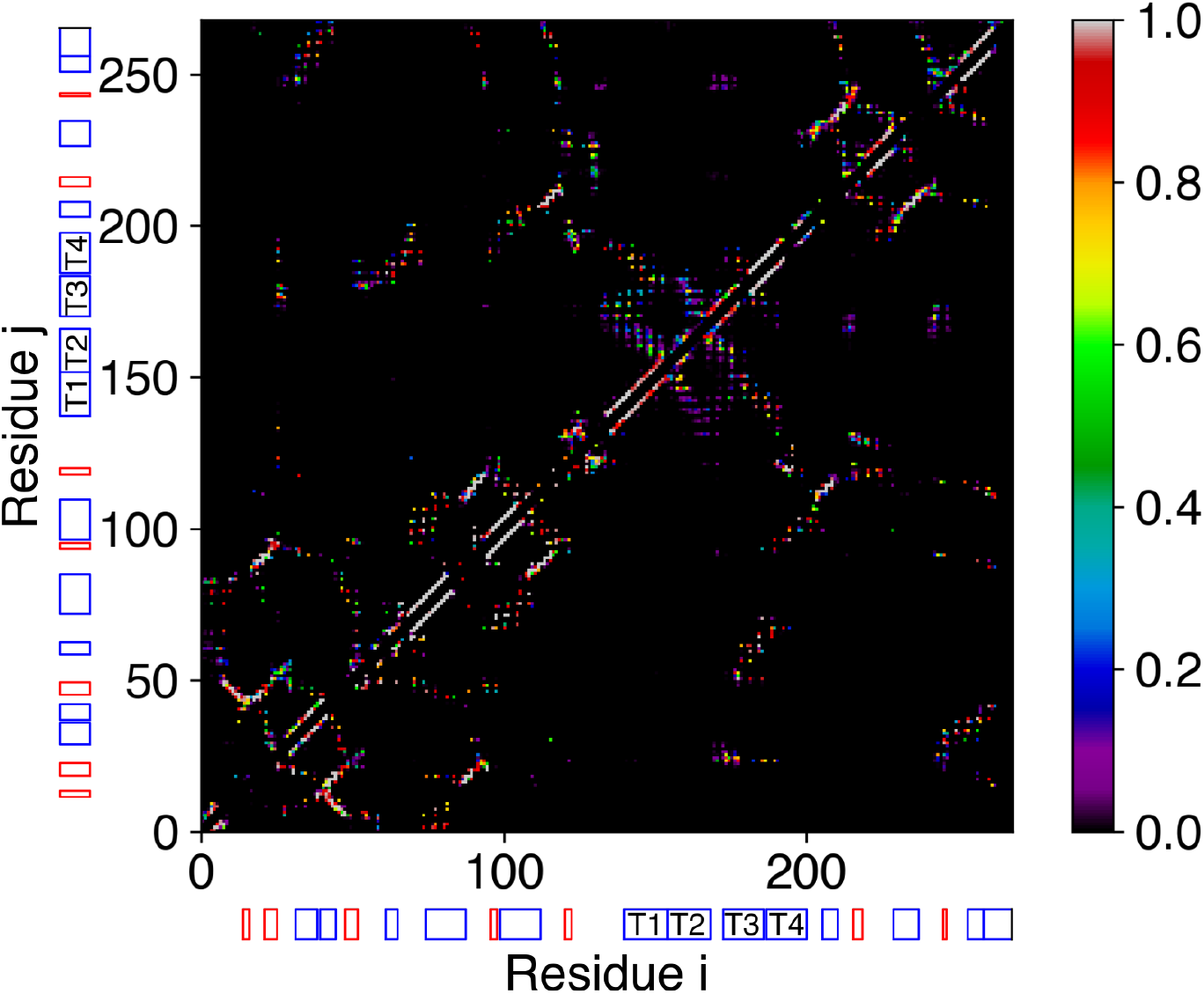
Contact probability matrices from kinetic Monte Carlo simulations of *At*D14 (bottom triangle) and *Sh*HTL7 (top triangle). Key secondary structure features are depicted by the red and blue rectangles.

**Fig. 5.**
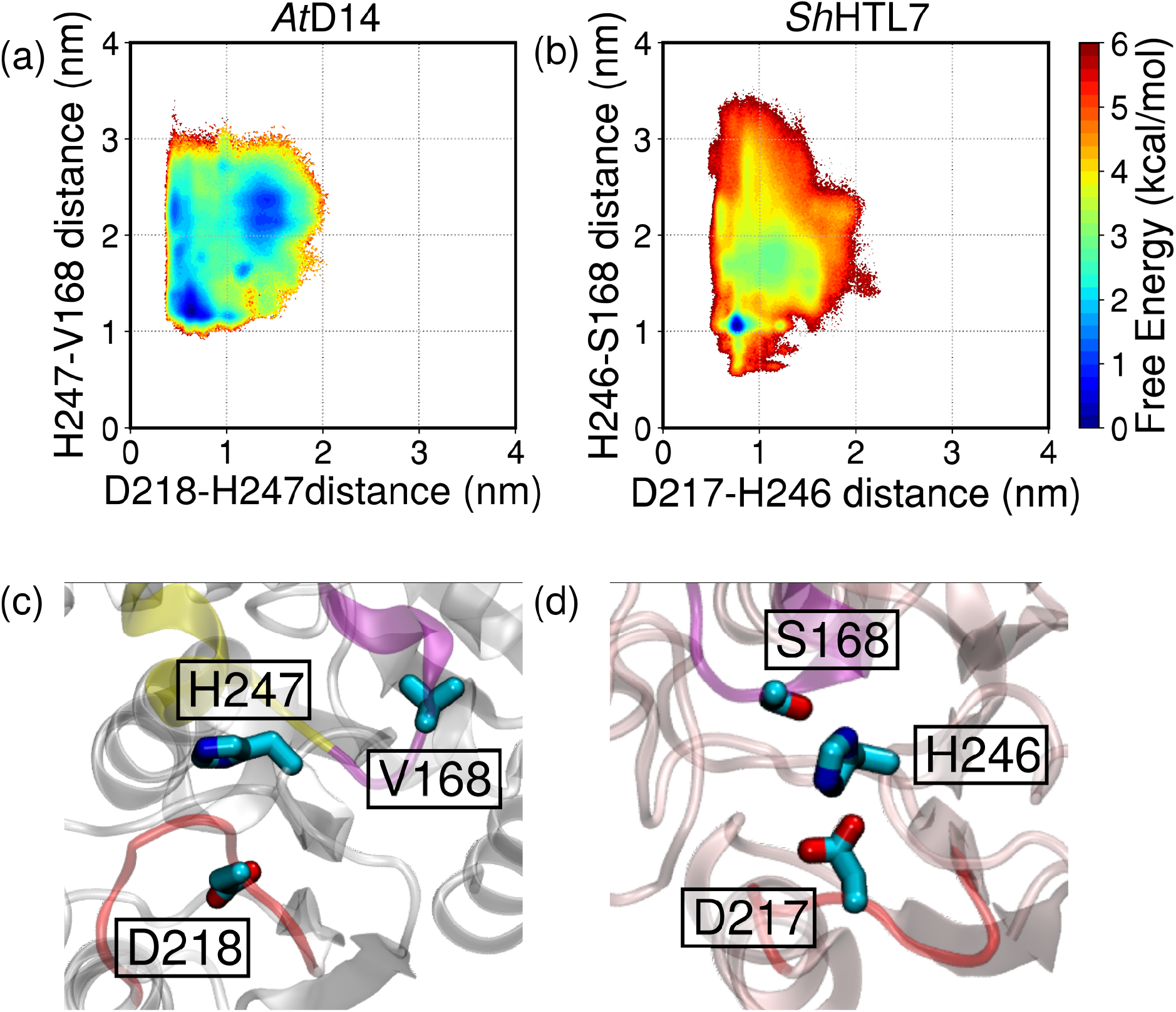
Free energy landscapes showing the coupling between the H246-S168 interaction and D217-H246 interaction in *Sh*HTL7 (b) and corresponding residues in *At*D14. An additional region of *Sh*HTL7 the landscape is seen where both contacts are formed which is not seen in *At*D14. Locations of the residues are shown in (c) for *At*D14 and (d) for *Sh*HTL7.

**Fig. 6.**
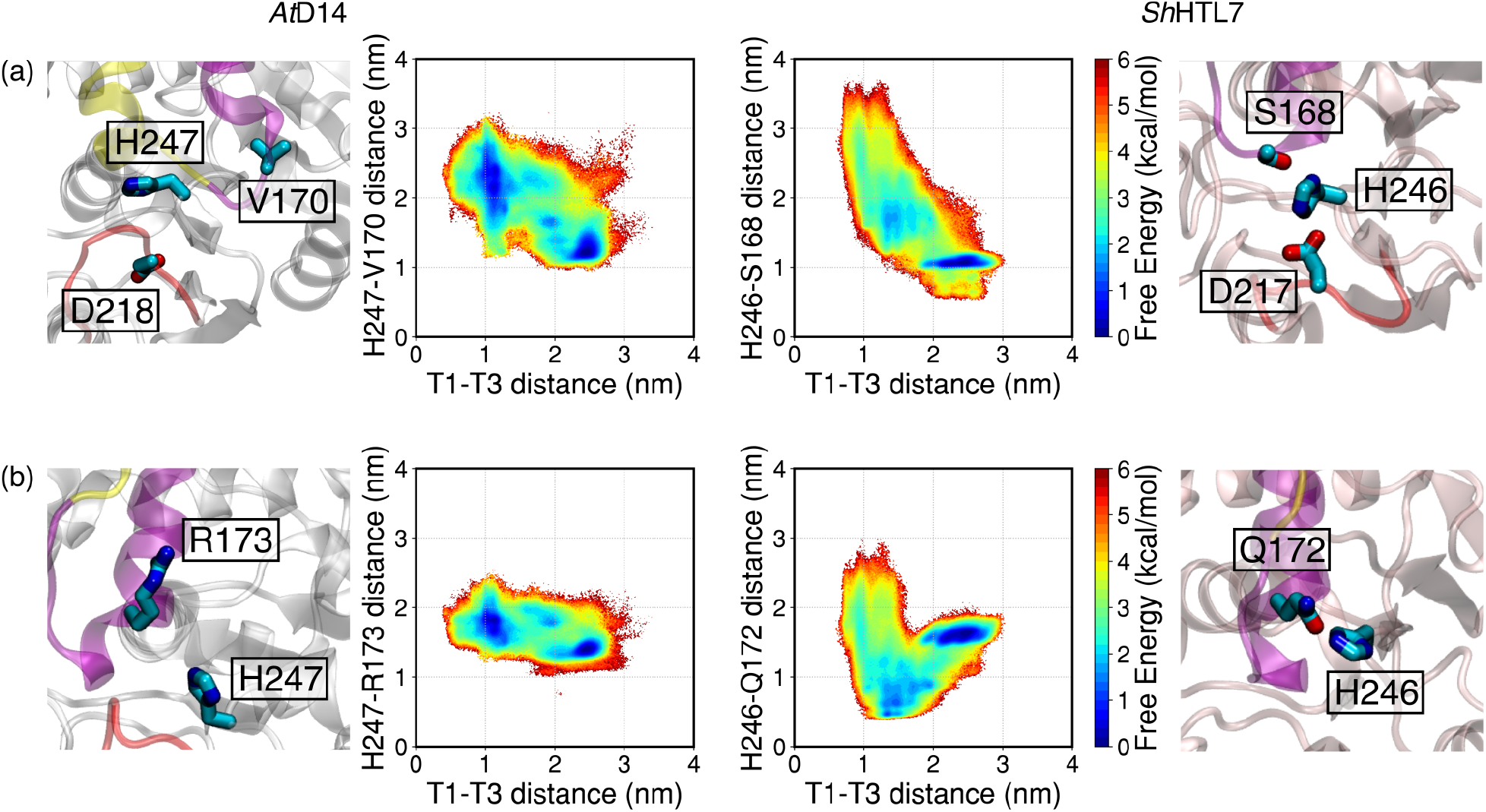
Key H247/246 interactions and their effect on T1-T3 helix distance. The presence of (a) S168-H246 and (b) Q172-H246 interactions in *Sh*HTL7 stabilize states in which the T3 helix is far from T1 and thus reduces the likelihood of T1-T3 helix closure.

The S168 residue on the T2-T3 loop in *Sh*HTL7 is able to form a hydrogen bond with H246 while the protein is in an active-like conformation which positions H246 so that it can interact with D217.

This prevents the formation of the full active state. This can be seen on the free energy landscape shown in Fig. 5, where the H246-D217 interaction is formed only when a S168-H246 interaction is formed. In contrast, the T2-T3 loop in *At*D14 does not interact with the H247 residue, so the same serine-histidine interaction that facilitates a D218-H247 interaction is absent. Thus, the D218 and H247 have a higher likelihood of being far apart in *At*D14 than in *Sh*HTL7. This increases the probability that the D-loop detaches, which in turn increases the probability of activation. Analysis of sequence conservation at this site shows again high variability among *At*D14 homologs with the most common residues being arginine, glutamine, and aspartate. In contrast, the site is a highly conserved serine among homologs of both *Sh*HTL7 and *At*KAI2. (Fig. S5).

The S168-H246 interaction also plays a role in preventing the T1 and T3 helices in *Sh*HTL7 from fully closing. The S168 residue is located close to the end of the T3 helix, thus its interaction with H247 allows it to access additional states in which the T1 and T3 helices are far apart which are not accessible to *At*D14 (Fig. 6a). This means that the lack of interaction between the T2-T3 loop and H247 in *At*D14 effectively removes a barrier to the T1-T3 helix closure motion, which increases the likelihood of T1-T3 helix closure and increases active state population.

Finally, the R173 residue in *At*D14 has a repulsive interaction with H247, while the equivalent Q172 residue in *Sh*HTL7 interacts with H246 more readily. Formation of this Q172-H246 interaction traps the protein in a state in which the T1 and T3 are partially closed, preventing full T1-T3 closure and thus full activation of the receptor. The absence of this interaction in *At*D14 prevents the formation of this off-pathway intermediate, thus lowering the barrier for full receptor activation. Analysis of conservation at this site again shows high variability of the site among close *At*D14 homologs, with most common residues at the site being glutamate and glutamine. The site is a highly conserved glutamine among close homologs of both *Sh*HTL7 and *At*KAI2.

## 3 Discussion

Using long time-scale molecular dynamics simulations, we have captured the activation processes of *At*D14 and *Sh*HTL7 in their *apo* forms. Our results indicate that *At*D14 has a higher active state population than *Sh*HTL7, which means that *At*D14 is more prone to *apo* activation than *Sh*HTL7. While this initially appears to contradict the idea that *Sh*HTL7 exhibits higher signaling activity than *At*D14, it is important to note that these measures of signaling activity were in the presence of a GR24 ligand. From an evolutionary perspective, it is possible that *Sh*HTL7 is less prone to activation in the absence of substrate because strigolactone from host plants acts as a germination signal, thus signaling activity would result in germination in the absence of a host plant. Additionally, *Sh*HTL7 has high binding affinity for MAX2, so *apo* receptor activation is highly likely to lead to a germination response. The evolution of parasitic plants involves weakened photosynthetic apparatus, which means germination in the absence of host plants would lead to the death of the parasite^35^. In fact, one widely proposed method of witchweed control is suicidal germination, which involves using small molecules to activate the strigolactone signaling pathway and induce germination in the absence of host crops^36^.

Using conditional probabilities of molecular switches, we have also found that D-loop detachment and pocket closure via merging of the T1 and T3 helices are the motions that distinguish between the activation pathways of *At*D14 and *Sh*HTL7. These can also be mechanisms to prevent *apo* activation. Previous studies indicate that the mutation of any catalytic triad residues prevents enzymatic activity. Thus, detachment of the D-loop prior to full activation would prevent catalytic activity even when all other molecular switches for activation are off, and the protein is in an inactive-like conformation. The ability to retain an enzymatically competent conformation prior to activation is likely poses an evolutionary advantage as well. While the necessity of strigolactone hydrolysis to formation of the active state of strigolactone receptors is disputed^7^, evidence exists that covalent modification of the catalytic histidine via hydrolysis of substrate is a driver of signaling^6,10^. If covalent modification of histidine is a driver of activation, higher population of inactive and catalytically compentent forms of the receptor would result in enhanced signaling activity in the presence of strigolactone hormones.

Finally, we identified several contact differences with the catalytic histidine residue (H247/246) that modulate differences in D-loop detachment and T1-T3 helix closure motions that are necessary for activation. Specifically, these contacts with H247/H246 decrease the likelihood of D-loop detachment and T1-T3 closure in *apo Sh*HTL7 to a larger degree than they do in *apo At*D14, thus making the relative population of the active conformation lower in *Sh*HTL7 than in *At*D14. Covalent modification to H247/246 would disrupt these interactions in both *At*D14 and *Sh*HTL7, however, since contacts with the catalytic histidine inhibit activation to a greater degree in *Sh*HTL7 than in *At*D14, modification to the histidine is likely to have a greater ability to enhance activation in *Sh*HTL7 than in *At*D14. Thus, covalent modification of histidine is likely to enhance association with MAX2 to a greater degree in *Sh*HTL7 than in *At*D14. This is consistent observation that *Sh*HTL7 shows high affinity for MAX2 and exhibits high signaling activity in the presence of GR24.

Due to the immense crop losses caused by *Striga* parasites, it is important to gain a mechanistic understanding of the strigolactone signaling process so that novel control strategies can be developed. Our work provides a step in understanding how the strigolactone receptors in *At*D14 and *Sh*HTL7 undergo the conformational transition from the inactive state to the active state that is able to associate with MAX2 proteins to induce downstream signaling effects. We also provide insight into a possible mechanism by which *Striga* parasites may evade suicidal germination in the absence of host plants. This provides a mechanistic target within *Sh*HTL7 by which to design suicidal germination stimulants or strigolactone signaling inhibitors to control witchweed.

## 4 Methods

### Molecular dynamics simulations

MD simulations were prepared using AmberTools 14/18 and run using Amber 14/18^37^. The protein was described using the ff14SB force field and water was described using the TIP3P model^38^. Initial structures for inactive *At*D14 and *Sh*HTL7 were obtained from Protein Data Bank entries 4IH4^39^ and 5Z7Y^40^, respectively. Active state models were created using the Modeller package with 5HZG as a template structure. Each system complex was solvated in a TIP3P water box of size ∼70 × 70 × 70 Å. NaCl was added at a concentration of 0.15M to neutralize the system. Each structure was minimized for 10000 steps using the conjugate gradient descent method and equilibrated for 10 ns. Production runs were performed for an aggregate of 457 *µ*s for *Sh*HTL7 and 249 *µ*s for *At*D14. Temperatures were held constant at 300 K using the Berendsen thermostat, and pressures were held constant at 1.0 bar using the Berendsen barostat. Full electrostatics were computed using the Particle Mesh Ewald algorithm with a cutoff distance of 10 Å ^41^. Bonds to hydrogen were constrained using the SHAKE algorithm^42^.

### Markov state model construction

Markov state models (MSMs) were constructed using the PyEmma^43^ package. 20 input distance features were computed using MDTraj 1.9.0^44^ (Table S2). The input distances were projected onto a reduced set of coordinates using time-lagged independent component analysis (TICA). The dimensionality-reduced coordinates were then clustered into states using the Mini-Batch K-Means algorithm prior to MSM estimation. The hyperparameters (number of TICA dimensions and number of clusters) were chosen via maximization of cross-validation scores (Fig. S3)^45^. Lag time was chosen by convergence implied timescales with respect to lag time (Fig. S2). Final parameters for MSM construction are listed in Table S3. Markovianity of the model was further validated using the Chapman-Kolmogorov test (Fig. S4). Free energy landscapes were calculated by computing the probability distribution along chosen sets of order parameters (*x, y*) and weighting each point by the equilibrium probability of its associated MSM state (Eq. 1).

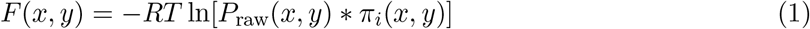

### Kinetic Monte Carlo simulations from MSMs

Kinetic Monte Carlo trajectories were generated from the MSM transition matrices by picking an initial state and appending further states by generating a random number *α* between 0 and 1 and choosing the corresponding state *S* as defined by Eq. 2.

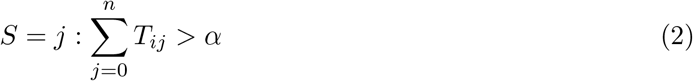

### Conditional probabilities of molecular switches

Conditional probabilities of the different molecular switches were calculated by computing using the definition of conditional probability (Eq. 3):

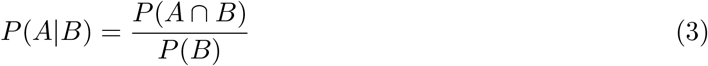

Eq. 3 can be used to derive a conditional probability for multiple conditions (Eq. 4).

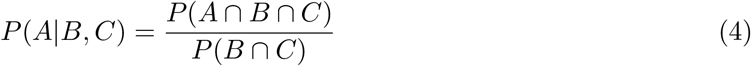

Probabilities of molecular switches were calculated by computing the probability of each molecular switch within in each MSM state, multiplying by respective MSM equilibrium probabilities for each MSM state, and summing over MSM states (Eq. 5):

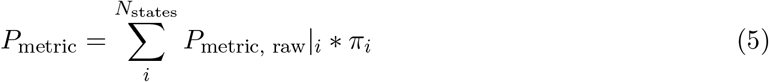

### Site conservation analysis

Sequence analysis was performed using the Consurf server^46,47^. For each protein, 150 closest homologs with sequence identity between 50% and 95% were selected from the UNIREF90 database. Full parameters are listed in Table S4.

## Supporting information

Supplementary Figures, Tables and References

