## Supplementary Figures, Tables and References for "Activation Mechanism of Strigolactone Receptors And Its Impact On Ligand Selectivity Between Host And Parasitic Plants"

### 1 Free energy landscapes projected onto TICA coordinates

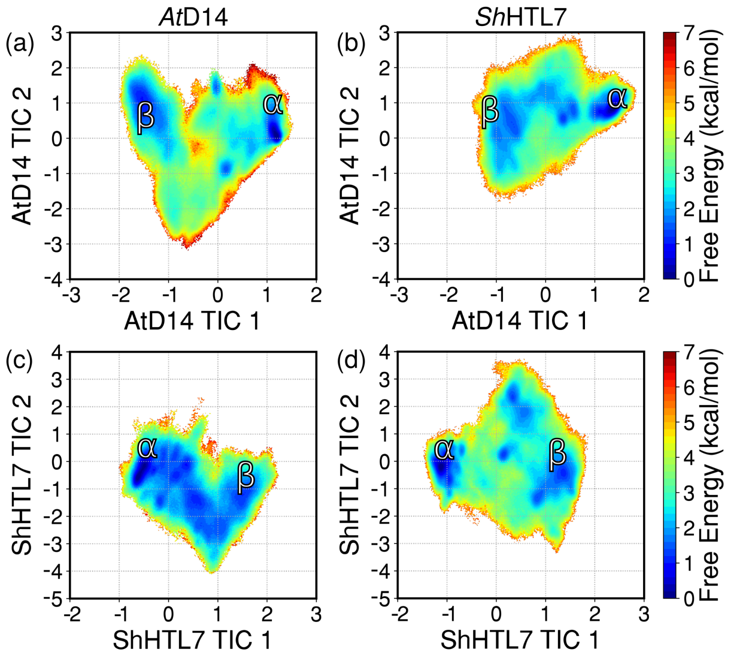


**Fig. S1.** Free energy landscapes of (a) *At*D14 activation projected onto *At*D14 TICA coordinates, (b) *Sh*HTL7 activation projected onto *At*D14 TICA coordinates, (c) *At*D14 activation projected onto *Sh*HTL7 TICA coordinates, (d) *Sh*HTL7 activation projected onto *Sh*HTL7 TICA coordinates. Inactive-like states are labeled as $\alpha$, and active-like states are labeled as $\beta$.

### 2 Conditional probabilities of molecular switches

| *At*D14 | T1-T3 closed | T1 extended | T2 unfolded | D-loop out |
|---|---|---|---|---|
| Given T1-T3 closed, T1 extended | | | 0.977 | 0.870 |
| Given T1-T3 closed, T2 unfolded | | 0.979 | | 0.866 |
| Given T1-T3 closed, D-loop out | | 0.956 | 0.950 | |
| Given T1 extended, T2 unfolded | 0.911 | | | 0.870 |
| Given T1 extended, D-loop out | 0.928 | | 0.979 | |
| Given T2 unfolded, D-loop out | 0.887 | 0.942 | | |
| *Sh*HTL7 | T1-T3 closed | T1 extended | T2 unfolded | D-loop out |
| Given T1-T3 closed, T1 extended | | | 0.917 | 0.639 |
| Given T1-T3 closed, T2 unfolded | | 0.925 | | 0.642 |
| Given T1-T3 closed, D-loop out | | 0.910 | 0.905 | |
| Given T1 extended, T2 unfolded | 0.362 | | | 0.658 |
| Given T1 extended, D-loop out | 0.367 | | 0.957 | |
| Given T2 unfolded, D-loop out | 0.305 | 0.800 | | |

**Table S1.** Conditional probabilities for each molecular switch given the presence of each pair of other molecular switch in *apo* *At*D14 and *Sh*HTL7. Each column of the table indicates the probabilities for each molecular switch, and each row corresponds to a different condition.

### 3 Distances used in MSM calculations

| | *At*D14 | *Sh*HTL7 |
|---|---|---|
| T1-T3 distances | A141-A171 | M139-S168 |
| | V144-S176 | T142-C173 |
| | M148-F180 | L146-F177 |
| T1-T2 distances | M148-W155 | L146-L153 |
| | V144-F159 | T142-T157 |
| | E140-L162 | V138-L160 |
| T1-T2 hinge contact distances | E149-E153 | D147-K151 |
| | A150-A154 | E148-S152 |
| | N151-W155 | N149-L153 |
| T2 helical contact distances | G158-L162 | G156-L160 |
| | F159-A163 | T157-L161 |
| | A160-V164 | A158-L162 |
| | P161-G165 | P159-A163 |
| D-loop-catalytic H distances | A216-H247 | S216-H246 |
| | K217-H247 | N216-H246 |
| | D218-H247 | D217-H246 |
| | V219-H247 | I218-H246 |
| | S220-H247 | M219-H246 |
| | V221-H247 | V220-H246 |
| | P222-H247 | P221-H246 |

**Table S2.** Featurizations used for MSM construction. C-$\alpha$ distances were used for all inter-residue distances.

### 4 Implied timescale plots used to select MSM lag times

MSM lag times were chosen using convergence of implied timescales. A lag time of 25 ns was chosen for both *At*D14 and *Sh*HTL7.

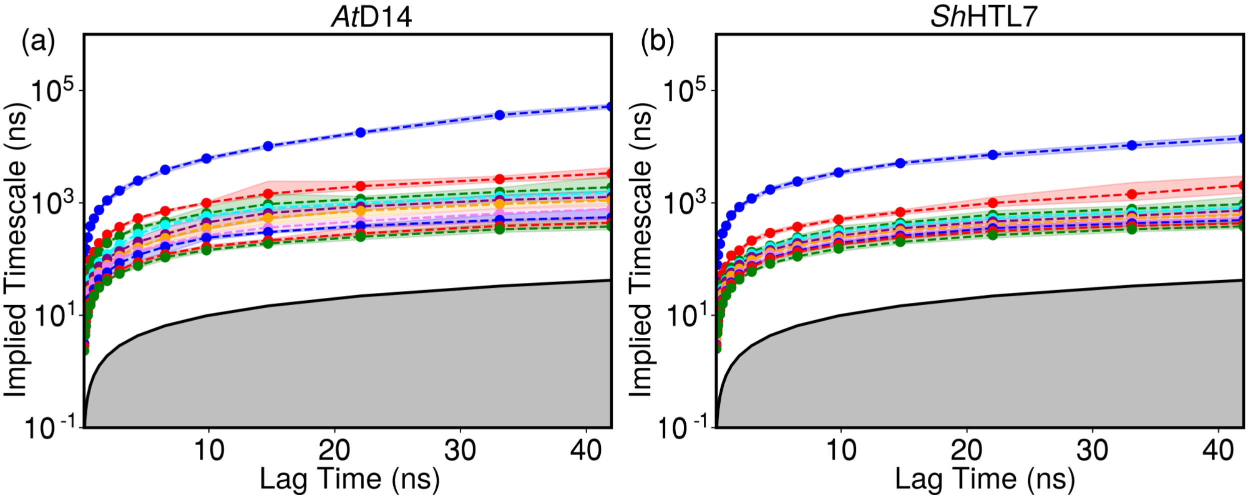


**Fig. S2.** Implied timescales of MSMs calculated at different lag times.

### 5 Results of cross-validation tests used to selected MSM hyperparameters

MSM hyperparameters were selected using GMRQ scores. Scores were calculated using shuffle-split cross validation with number of TICA components ranging from 2 to 8 and cluster numbers ranging from 100 to 500. Selected parameters are shown in Table S3.

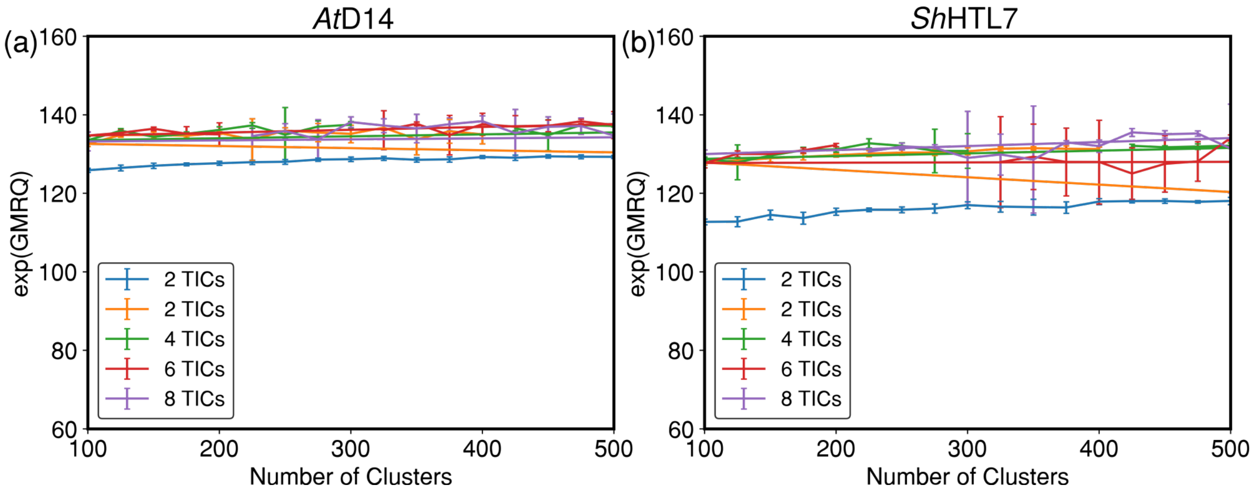


**Fig. S3.** Cross-validation scores with varying numbers of TICA components and clusters

| | *At*D14 | *Sh*HTL7 |
|---|---|---|
| Lag time (ns) | 25 | 25 |
| Number of TICA components | 10 | 8 |
| Number of clusters | 400 | 275 |

**Table S3.** Final parameters used for MSM construction

### 6 Chapman-Kolmogorov validation of MSMs

Validation of MSMs was performed using the Chapman-Kolmogorov test. Briefly, this tests for Markovianity of a system by comparing estimated transition matrices at lag time $n\tau$ with predictions of the same transition matrix by multiplying a transition matrix calculated using lag time $\tau$ $n$ times.

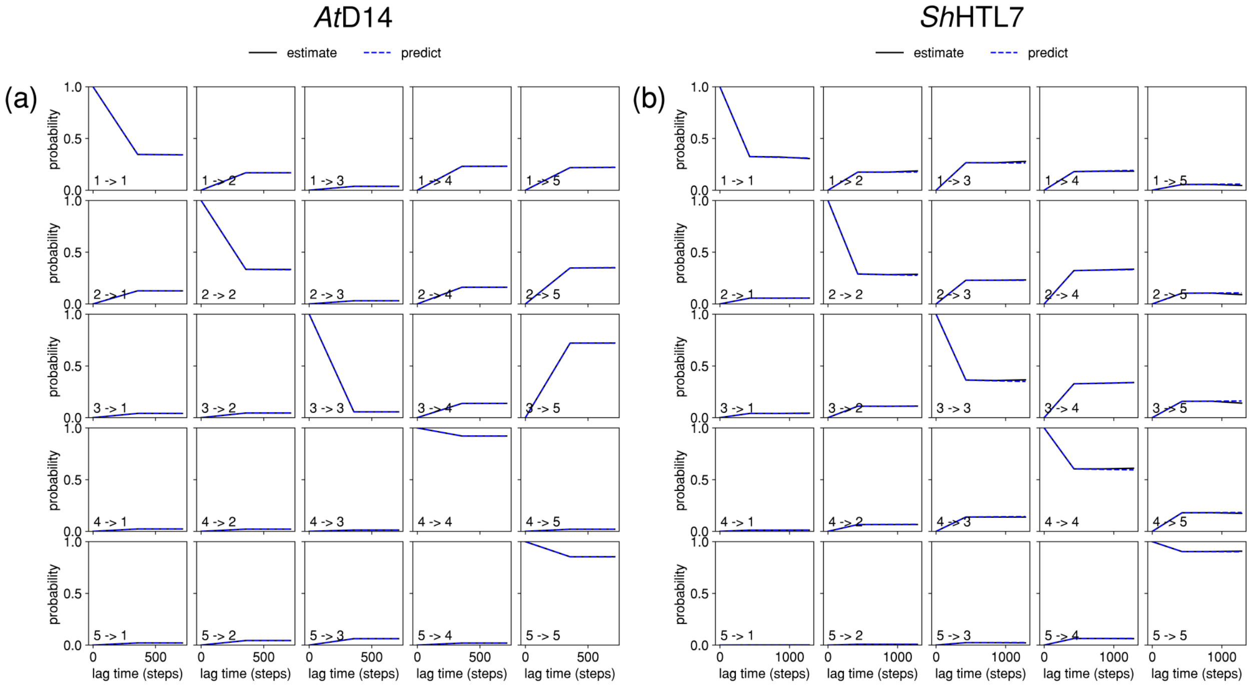


**Fig. S4.** Chapman-Kolmogorov validation for *At*D14 and *Sh*HTL7 MSMs.

### 7 Site conservation of key residues

| | *At*D14 | *Sh*HTL7 | *At*KAI2 |
|---|---|---|---|
| PDB ID | 4IH4 | 5Z7Y | 4HRX |
| Multiple Sequence Alignment | MAFFT | MAFFT | MAFFT |
| Homolog Source | UNIREF90 | UNIREF90 | UNIREF90 |
| Homolog Search Algorithm | HMMER | HMMER | HMMER |
| HMMER E-value | 0.0001 | 0.0001 | 0.0001 |
| HMMER Iterations | 1 | 1 | 1 |
| Maximal % Identity | 95 | 95 | 95 |
| Minimal % Identity | 50 | 50 | 50 |
| Number of Sequences | 150 | 150 | 150 |

**Table S4.** Parameters used for calculation of site conservation

| Site | Conservation Score | Most Frequent Residues |
|---|---|---|
| *At*D14 Homologs | | |
| A216 | 0.836 | T,F,S,D,H,V,M,K,L,Q,G,N,A,R,E |
| K217 | 1.090 | K,T,Y,S,D,R,E,G,N,I,V,M,P,Q,H,A |
| S220 | -0.716 | V,M,L,A,F,I,S |
| V168 | -0.016 | R,E,N,A,M,V,K,P,L,Q,F,T,D,S,H |
| R173 | 1.436 | K,Q,M,V,H,S,D,T,E,R,I,N,G,A |
| *Sh*HTL7 Homologs | | |
| S215 | 1.226 | G,K,C,L,T,R,I,A,E,M,X,S,V |
| N216 | -0.238 | K,T,A,R,N,M,Q,E |
| M219 | 0.162 | G,K,C,L,T,R,I,A,E,M,X,S,V |
| S168 | -0.923 | T,S |
| Q172 | -1.035 | Q |
| *At*KAI2 Homologs | | |
| V215 | 1.612 | V,F,M,R,G,A,L,T,K,S,E,N,I |
| K216 | -0.089 | Y,N,K,R,T,M,A |
| A219 | -0.732 | A,V,S,I |
| S168 | -0.753 | A,T,S,C |
| Q172 | -0.935 | Q |

**Table S5.** Conservation of key sites in *At*D14, *Sh*HTL7, and *At*KAI2 and most frequent residues.

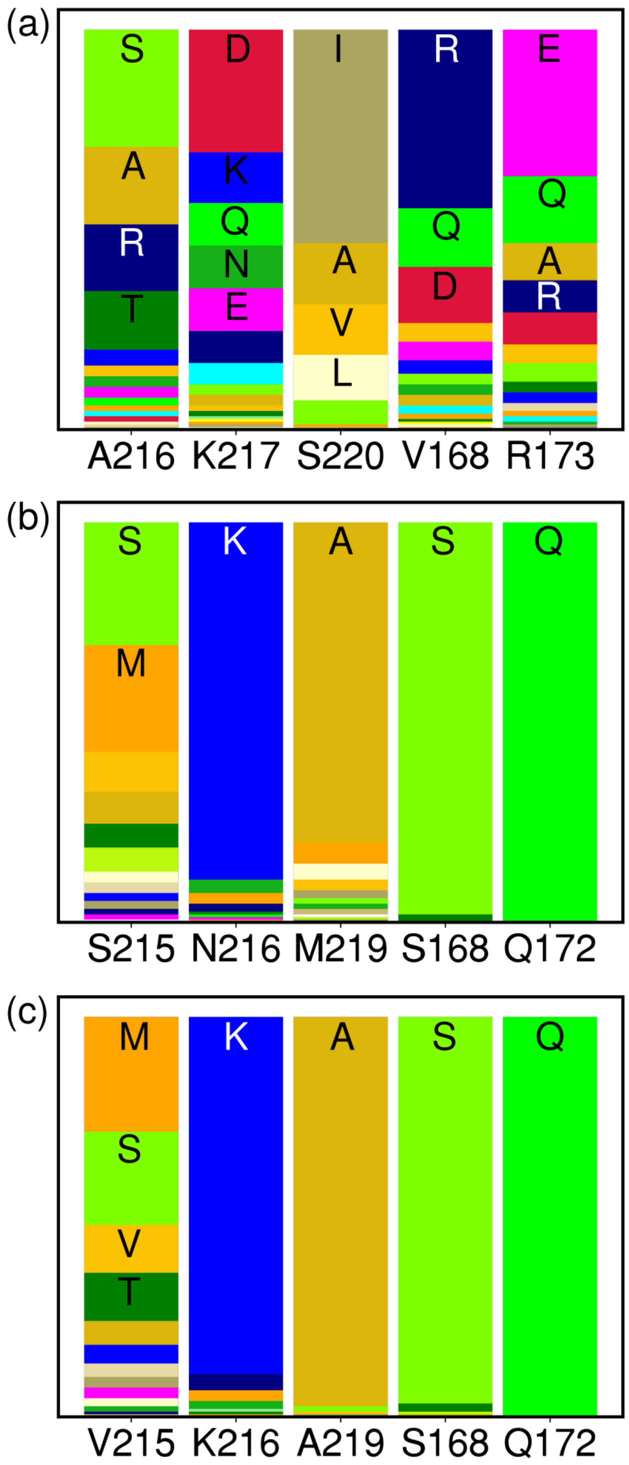


**Fig. S5.** Most commonly occurring residues at key sites among close homologs of (a) *At*D14, (b) *Sh*HTL7, and (c) *At*KAI2.

### 8 Contacts with high contact probability difference vs D-loop distance

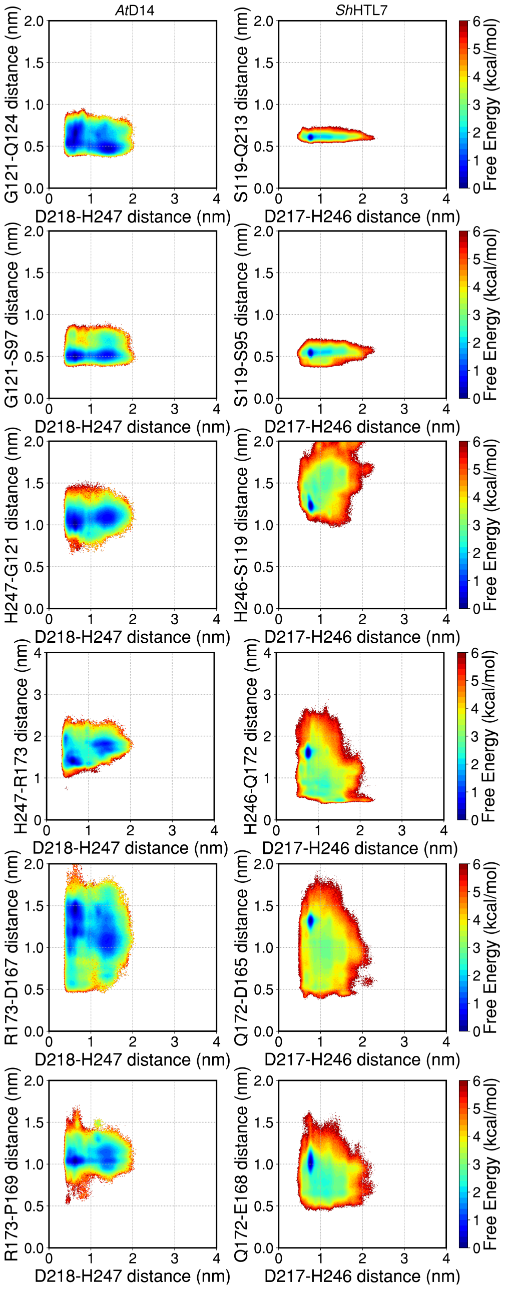


**Fig. S6.** Free energy landscapes projected onto catalytic D-H distance and distances with high difference in probability between *At*D14 and *Sh*HTL7

### 9 Contacts with high contact probability difference vs T1-T3 distance

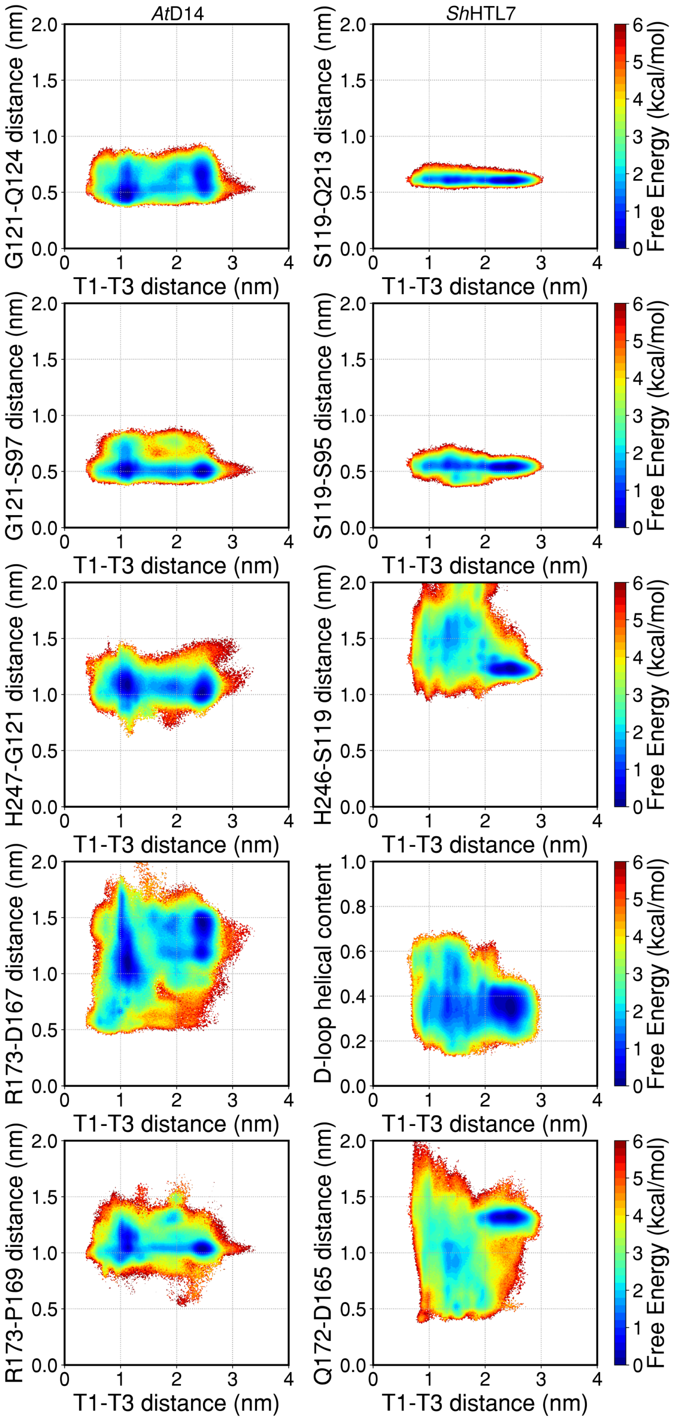


**Fig. S7.** Free energy landscapes projected onto catalytic D-H distance and distances with high difference in probability between *At*D14 and *Sh*HTL7
